# Study of an *FBXO7* patient mutation reveals Fbxo7 and PI31 co-regulate proteasomes and mitochondria

**DOI:** 10.1101/2021.12.22.473884

**Authors:** Sara Al Rawi, Lorna Simpson, Neil Q. McDonald, Veronika Chernuha, Orly Elpeleg, Massimo Zeviani, Roger A. Barker, Ronen Spiegel, Heike Laman

## Abstract

Mutations in *FBXO7* have been discovered associated with an atypical parkinsonism. We report here a new homozygous missense mutation in a paediatric patient that causes an L250P substitution in the dimerization domain of Fbxo7. This alteration selectively ablates the Fbxo7-PI31 interaction and causes a significant reduction in Fbxo7 and PI31 levels in patient cells. Consistent with their association with proteasomes, L250P patient fibroblasts have reduced proteasome activity and proteasome subunits. We also show PI31 interacts directly with the MiD49/51 fission adaptor proteins, and unexpectedly, PI31 acts as an adaptor enabling SCF^Fbxo7^ ligase to ubiquitinate MiD49. Thus, the L250P mutation changes the function of Fbxo7 by altering its substrate repertoire. Although MiD49/51 expression was reduced in L250P patient cells, there was no effect on the mitochondrial network. However, patient cells had higher levels of ROS and reduced viability under stress. Our study shows that Fbxo7 and PI31 affect each other’s functions in regulating both proteasomal and mitochondrial function and demonstrate a new function for PI31, as an adaptor for the SCF^Fbxo7^ E3 ubiquitin ligase.

## Introduction

Parkinson’s disease (PD) is the second most common neurodegenerative disorder in the world, affecting more than 6 million people worldwide, where 90% of cases are idiopathic. Over the past two decades, an increasing number of mutations have been identified and associated with familial forms of PD. Study of the functions of these so-called PARK genes provides new opportunities to understand the mechanisms underlying familial and sporadic forms of Parkinson’s disease (1, 2). Since 2008, when the first case of Parkinsonian pyramidal syndrome in an Iranian family was linked to a mutation in *FBXO7*, many other pathological recessive mutations of *FBXO7* have been identified (3–6). These mutations are clustered at the N- and C-terminal domains of Fbxo7, and a wide range of symptoms have been reported (2, 6–16). The age of the onset is also variable with cases arising in childhood and adolescence to young adults and older patient. Some patients, for example, with L34R mutations have a disease-onset and presentation similar to idiopathic cases of Parkinson’s disease (6).

Fbxo7 was originally identified as a substrate recognition subunit of Skp1-Cullin-1-F-box (SCF)-type E3 ubiquitin ligases. It has a C-terminal proline-rich (PRR) domain and an N-terminal ubiquitin-like (Ubl) domain, which are involved in recruiting substrates for ubiquitination by the SCF^Fbxo7^ ligase (17). Although hundreds of substrates of SCF^Fbxo7^ ligase have been identified, many are not subject to proteasomal degradation (18). Moreover, some Fbxo7 interacting proteins are not ubiquitinated, but rather scaffolded and stabilised by association with Fbxo7. Examples include cell cycle regulators, Cdk6 and p27, and consequently, Fbxo7 augments G1 phase cyclin D/Cdk6 activity (19–21). Fbxo7 also regulates the levels of PINK1/PARK6, another autosomal recessive gene associated with Parkinson’s disease (22). Fbxo7 acts in a common pathway with PINK1 and Parkin/PARK2 to enable stress-activated mitophagy, and it also affects basal mitochondrial function (23, 24).

In addition to two substrate recruiting domains, Fbxo7 also contains an FP (Fbxo7-PI31) dimerization domain that enables its homo-dimerization and hetero-dimerization with the proteasome regulator, PI31/PSMF1 (25, 26). Fbxo7 and PI31 share a similar organisational structure; both contain an FP domain and a PRR domain (26). In many cell types, a reduction in Fbxo7 levels correlates with lower PI31 levels (19, 27, 28), indicating Fbxo7 also stabilises PI31.

Multiple global proteomic profiling and interactome studies have demonstrated the association of Fbxo7 and PI31 with each other and the proteasome (29–32). The PRR domain of PI31 mediates its interaction with α subunits of the 20S proteasome, although the effect of PI31 on proteasome activity is controversial (33). Fbxo7, has been reported to ubiquitinate a proteasome subunit α2/PSMA2, regulating proteasome assembly (34). A mouse with conditional expression of Fbxo7 in motor neurons showed early-onset motor deficits and premature death which was attributed to reduced proteasome activity (34). More recent work has detailed a role for Fbxo7 in association with PI31 in the axonal transport of proteasomes (35). These studies suggest a multi-faceted role for Fbxo7 in the assembly, activity, and localisation of proteasomes in neurons.

Here we report a new homozygous point mutation in *FBXO7* in a paediatric patient from an Israeli family presenting with a combination of upper and lower motor neuronal problems and dystonia. At their initial clinical presentation at 5 months of age, whole exome sequencing was performed, and a novel point mutation c.749T>C in the *FBXO7* gene (p.Leu250Pro, NM_012179.4) was identified. We investigated the molecular effects of the L250P point mutation, located in the FP domain of Fbxo7, which selectively ablates its interaction with PI31. We found that the mutant Fbxo7-L250P and PI31 levels were significantly reduced in patient cells, which have decreased levels of proteasome subunits and proteasome activity. We then went on to show that PI31 acts as an adaptor for SCF^Fbxo7^ ligase and that MiD49 and MiD51, fission adaptor proteins, were direct interacting partners for PI31. Although MiD49 and MiD51 both interact with Fbxo7, PI31 enables only MiD49 ubiquitination by SCF^Fbxo7^ ligase. MiD49 and MiD51 levels were significantly reduced in L250P patient cells, although no effects on the dynamics of the mitochondrial network were evident. Finally, we found that the patient cells had a significantly reduced mitochondrial mass and higher levels of cellular and mitochondrial ROS, and reduced viability under stress. Our data reveal the dual functions of Fbxo7 and PI31 in regulating both proteasomal and mitochondrial function.

## Results

### Demographic data and detection of a new pathogenic FBXO7 point mutation (L250P)

A female patient, the first offspring of healthy related parents, initially presented at the age of 5 months with decreased limb movements and evolving truncal hypotonia. Her neurological examination revealed head drop and axial hypotonia with reduced facial and limb movements, generalized hyper-reflexia associated with bilateral ankle clonus and positive Babinski sign consistent with a pyramidal syndrome. Brain MR imaging at the age of 1.5 years showed thinning of corpus callosum, but otherwise was unremarkable.

Disease course was marked by the development of limb rigidity and fluctuating dystonic postures, but cognitive and communication skills remained relatively preserved. Ophthalmological examination at the age of 2.5 years displayed bilateral optic pallor with decreased visual evoked potential responses consistent with an evolving optic atrophy.

Single whole exome sequencing (WES) analysis of her DNA, extracted from whole blood, identified a homozygous c.749T>C in the *FBXO7* gene (p.Leu250Pro, NM_012179.4). This variant was predicted to be pathogenic by web based applications that perform *in silico* tests including Mutation Taster (http://mutationtaster.org), Polyphen 2 (http://genetics.bwh.harvard.edu/pph2) and SIFT (http://siftdna.org).

Given the neurological phenotype of the child dominated by axial hypotonia, limb rigidity, decreased spontaneous movements and generalized pyramidal signs along with a homozygous variant in *FBXO7*, a gene known to be associated with parkinsonian pyramidal syndrome, a dopamine transporter (DAT) SPECT imaging was performed which showed lack of uptake in the striatum. She started treatment with L-DOPA with gradual dose increase. She showed an initial good response with an increased range of limb movement, but the long-term clinical effect waned. In addition, she developed progressive kyphoscoliosis that is related to her rigidity and urinary retention that necessitated regular catheterization. A repeated brain MRI at 3.5 years was similar to her initial imaging. On her last examination at 5 years of age, the child expresses very slow but steady progress in her developmental milestones. She is able to sit with support, stand in a walker, grasp objects and bring them to her mouth. She utters two words and communicates with her family. However, in addition to her initial motor impairment, the patient developed moderate global developmental delay, and her recent developmental quotient (DQ) is 40.

### L250P point mutation localises to the FP domain of Fbxo7 and ablates Fbxo7 hetero-dimerization with PI31

The L250P mutation is located within the FP domain (residues 180-324) of Fbxo7, which enables its homo-dimerization and hetero-dimerization with and FP domain in PI31 (26). The FP domains of both are structurally similar adopting an α/β-fold with a central five-stranded anti-parallel β-sheet flanked by two α-helices in the N-terminal and three α-helices at the C-terminus (25, 26, 36). The FP domain uses an αβ-interface for FP-FP-domain interaction and dimerization, although different modalities for interacting have been proposed (αβ, αα or ββ—see ref (25)). For example, the Fbxo7 interface has been suggested to use the β-interface (Fig. 1A, lefthand panel, green surface). Leucine 250 is a buried hydrophobic core residue within the central beta strand of the Fbxo7 FP domain β-sheet. Substitution by proline (L250P) would be particularly disruptive to the β strand resulting in a bulge or kink to the strand. The kink arises as the proline sidechain prevents a hydrogen on the amide nitrogen, so it cannot act as a hydrogen-bond donor, thereby disrupting the mainchain hydrogen-bonding network of a sheet. Also, the proline sidechain places a steric constraint on the neighbouring residues because of its pyrrolidine ring, perturbing its secondary structure (Fig. 1A). Thus, the L250P mutation would be predicted to both partially destabilise the Fbxo7 FP domain as well as disrupt dimerization through its β-interface with the FP domain of PI31 (25, 26, 36).

**Figure 1:**
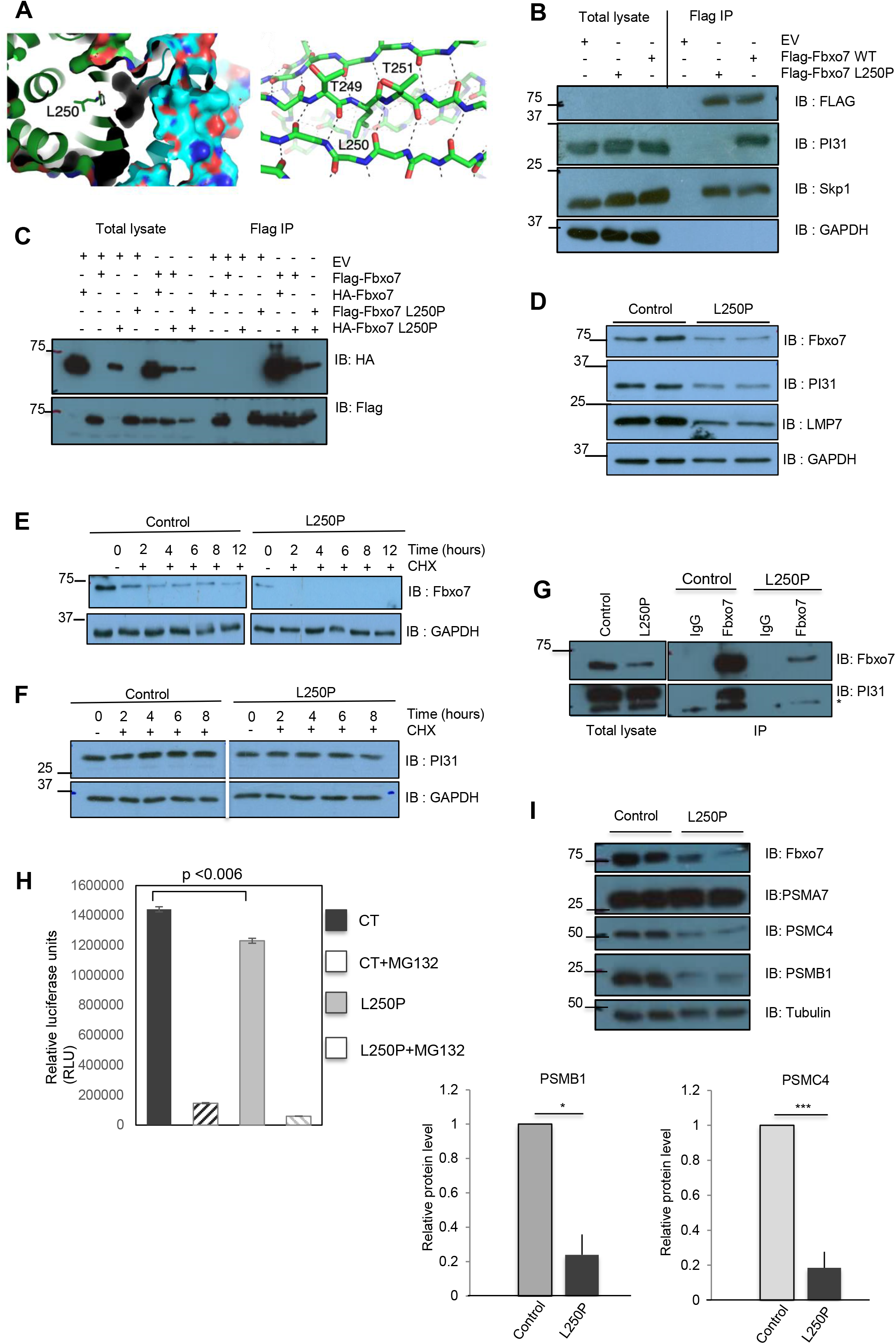
The FP domain missense mutation L250P compromises Fbxo7-PI31 interaction and proteasome activity. **(A)** Close-up of the structure of Fbxo7 FP domain highlighting the position of the L250 residue within the central beta strand of the Fbxo7 FP domain β-sheet. The Fbxo7 FP domain is on the left (green) and PI31 on the right (cyan α-interface). The β-interface is the proposed Fbxo7-PI31 heterodimerization surface. **(B)** Co-immunoprecipitation of endogenous PI31 with Flag-Fbxo7 WT or L250P in total cell lysates from 293T cells transfected with Flag-Fbxo7 WT or L250P. Immunoblots of total lysate prior to anti□FLAG immunoprecipitation are also shown (n=2). **(C)** Co-immunoprecipitation of Flag-Fbxo7 WT or L250 and HA-Fbxo7 WT or L250P in total cell lysates from 293T cells transfected with Flag-Fbxo7 WT or L250P and HA-Fbxo7 WT. Immunoblots of total lysate prior to anti□FLAG immunoprecipitation are also shown (n=3). **(D)** Immunoblotting for the expression of Fbxo7, PI31 and LMP7 proteins from cell lysates of control or L250P fibroblasts. GAPDH serves as a loading control (n=4). **(E, F)** Cycloheximide (CHX) chase on control and L250P fibroblasts. Total cell lysates were prepared after the indicated treatment time and the levels of Fbxo7 (**E**) and PI31 (**F**) were analysed by immunoblot (n=3). **(G)** Co□immunoprecipitation from control or L250P cells lysates. Lysates were immunoprecipitated with antibodies to Fbxo7 and immunoblotted for PI31 (n=1). **(H)** Proteasome activity assay performed on total cell lysates from control or L250P fibroblasts. MG132 treatment is used to confirm the efficiency of the assay by blocking the proteasome and diminishing the signal (n=2). **(I)** Immunoblot analysis for the proteasome subunits PSMA7, PSMC4 and PSMB1 in total cell lysates from control and L250P fibroblasts. Tubulin is used as a loading control (n=6).

To explore the effect of the L250P mutation on Fbxo7 dimerization with PI31, co-IP experiments were performed. HEK293T cells were transfected with plasmids expressing Flag-tagged WT or mutant Fbxo7. Anti-Flag immunoprecipitates of total cell lysates were assayed by immunoblotting. Endogenous PI31 associated with WT Fbxo7 but not L250P Fbxo7, indicating that the L250P mutation ablated their interaction (Fig. 1B). We also tested whether the L250P mutation affected Fbxo7 interaction with Skp1, which binds the F-box domain adjacent to the FP domain. Both alleles of Fbxo7 interacted with Skp1, indicating that only the interaction with PI31 is affected. Moreover, the ability of Fbxo7-L250P to bind Skp1 suggests it can be incorporated into an SCF-E3 ubiquitin ligase, although lacking PI31. These data show PI31 hetero-dimerizes with Fbxo7 exclusively via its β-interface and raises the possibility that PI31 acts as an adaptor for SCF^Fbxo7^ ligase.

Co-immunoprecipitation experiments were also conducted to test the effect of L250P mutation on Fbxo7 homo-dimerization through either an αβ or ββ interface. For these experiments HEK293T cells were transfected with FLAG-WT or L250P alleles of Fbxo7 along with HA-tagged WT or Fbxo7-L250P alleles. Anti-FLAG immunoprecipitates were assayed for the presence of WT or mutant Fbxo7-HA protein. We noted the Fbxo7 L250P-HA constructs were less well expressed compared to WT Fbxo7-HA constructs. Nonetheless, both WT and mutant Fbxo7 were detected in WT and mutant FLAG-immunoprecipitates (Fig. 1B), indicating the L250P mutation does not prevent Fbxo7 homo-dimerization. These data indicate Fbxo7 FP is not misfolded by the mutation and that homo-dimerization is not dependent exclusively on the β-interface of its FP domain and suggest alternate interfaces mediate this.

### Patient fibroblasts with L250P mutation show reduced expression and stability of Fbxo7 and PI31

We previously showed Fbxo7 binding to PI31 stabilises its levels in multiple cell types (19, 27, 28). Because this interaction is specifically ablated by the L250P mutation, we predicted PI31 expression levels would be reduced in patient fibroblasts. We assayed PI31 protein levels by immunoblotting total cell lysates from patient and control fibroblasts. Steady state PI31 levels were 45% lower in patient cells compared to the control (Fig. 1D). We also measured the effect of the L250P mutation on the expression levels and stability of Fbxo7 and found there was approximately a 50% reduction of Fbxo7-L250P protein compared to WT Fbxo7 (Fig 1D). To measure the protein half-life, we performed a time course of treatment with the cycloheximide. Fbxo7 was rapidly degraded in the patient fibroblasts with a half-life of less than 2 hours, compared to the control cells where Fbxo7 had a half-life of 2 hours (Fig. 1E). However, the half-life of PI31, was unaffected (Fig. 1F). These data demonstrate the reciprocal stabilisation of Fbxo7 and PI31 levels conveyed through their hetero-dimerization and suggest Fbxo7 and PI31 influence each other’s functions.

We confirmed the effect of the Fbxo7-L250P mutation on its interaction with PI31 in patient fibroblasts. To mitigate against the reduced expression of these proteins in patient cells, total cell lysates were prepared from 2.5 times the number of patient fibroblasts as control fibroblasts. Immunoprecipitation of the endogenous Fbxo7 in control cell lysates showed a robust interaction with PI31, but no interaction was detected in the patient cell lysates (Fig. 1G).

### Reduced levels of LMP7 and proteasome activity in Fbxo7 L250P patient fibroblasts

Fbxo7 and PI31 are found associated with proteasomes, and because their levels were significantly reduced in patient cells, we tested the effect on proteasome activity. We performed a chymotrypsin-like activity assay on lysates made from cells cultured with or without the proteasome inhibitor MG132. We detected a significant 15% decrease in proteasomal activity in patient fibroblasts compared to control fibroblasts (Fig. 1H). We tested whether the decrease in proteasome activity was due to reduced levels of proteasome subunits by immunoblotting lysates for PSMA7, PSMB1, and PSMC4 (Fig. 1I). We noted that PSMB1 and PSMC4 showed a significant reduction in levels (Fig. 1I); however, PSMA7 levels were similar in both cell lines. Since PI31 has additionally been described as a regulator of the stress-induced immunoproteasome, which contains alternate beta subunits, like LMP2, MECL-1, and LMP7 (37–39), cell lysates from patient cells were also immunoblotted for LMP7, which were also reduced (Fig. 1D). These data suggest reduced proteasome activity in the patient cells is due to lower expression of constitutive and alternate proteasome subunits.

### Fbxo7 and PI31 interacts with the mitochondrial fission adaptors MiD49 and MiD51

The Fbxo7 L250P mutation prevented PI31, but not Skp1 binding, which raised the possibility that PI31 acts as an adaptor for SCF^Fbxo7^ ligase. To identify SCF^Fbxo7^ substrates reliant on PI31, we cross-referenced the BioGRID database for PI31/PSMF1 interactors with our own experiments to identify SCF^Fbxo7^ substrates using protein arrays. A 2005 study reported PI31 and MiD49/SMCR7/MIEF2 as high confidence interacting proteins (32, 40), and we previously identified MiD49 as a lower confidence *in vitro* substrate of SCF^Fbxo7^ ligase (18). MiD49 and the closely related MiD51 protein are important fission adaptor proteins that recruit the Drp1 GTPase to organelles, including mitochondria, ER, lysosomes, undergoing fission (Fig. S1A) (41–43). To validate an interaction between the MiD49 and PI31, we first performed co-IP assays. MiD49-Myc was cotransfected with either FLAG-tagged WT PI31 or a PI31 mutant lacking the C-terminal proteasome-interacting domain (ΔC, aa1-189) in HEK293T cells expressing either a control shRNA or a shRNA targeting Fbxo7 expression. MiD49 was detected in anti-FLAG immunoprecipitates for both WT PI31 and PI31ΔC (Fig. 2A). PI31 binding to MiD49 was also detected in cells with undetectable Fbxo7 expression. The interaction between PI31 and MiD49 was also verified using *in vitro* pull-down assays where GST-PI31 fusion proteins were used to pull down MiD49 made using reticulocyte lysates (Fig 2B). Together these data indicate PI31 and MiD49 interact directly and independently of Fbxo7 and the C-terminus of PI31.

**Figure 2:**
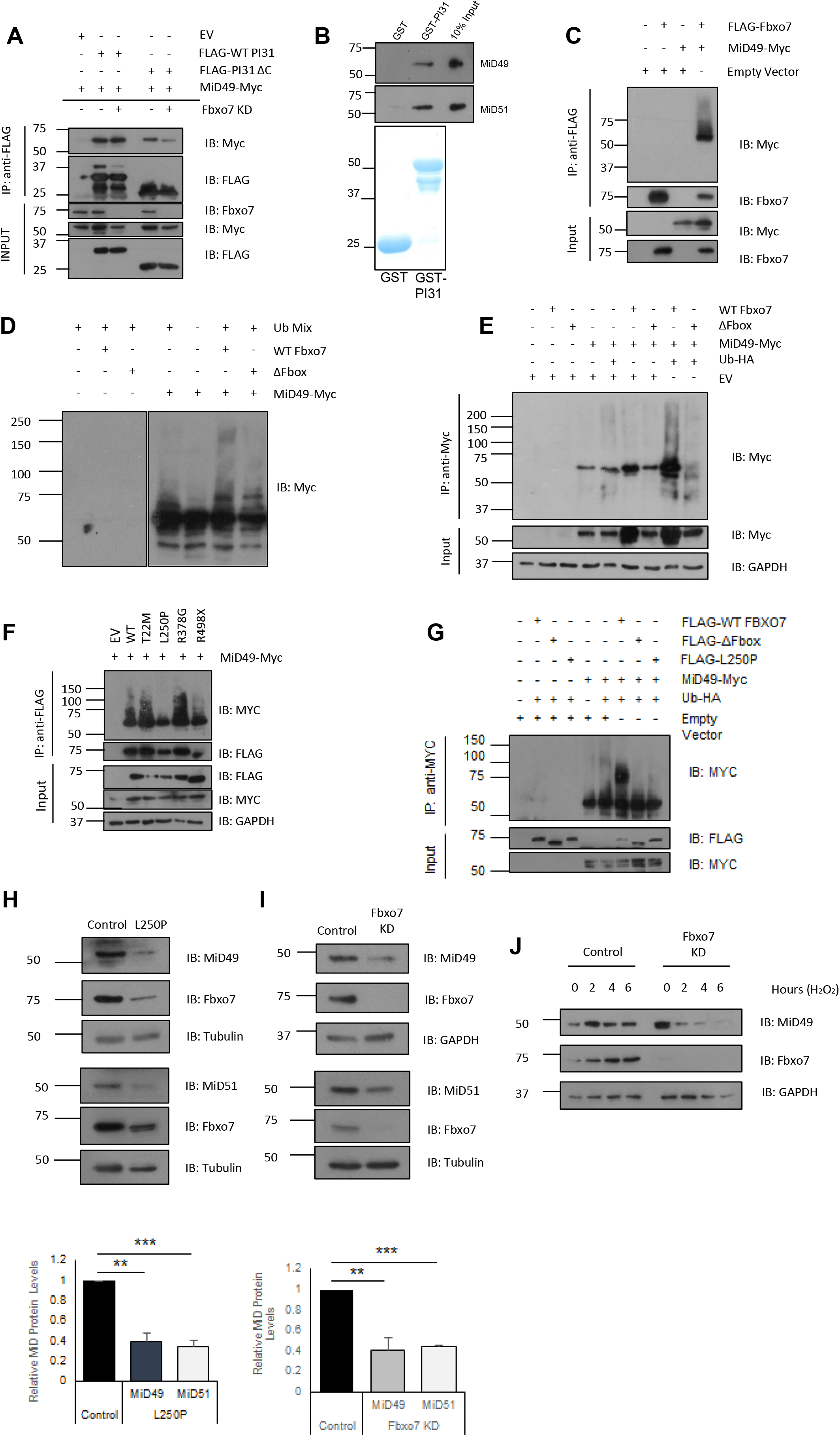
Fbxo7 and PI31 interact with the mitochondrial fission adaptors MiD49 and MiD51. **(A)** Co-immunoprecipitation of MiD49 and PI31 from control and Fbxo7 knock-down HEK293T cells co transfected with FLAG-tagged PI31 and Myc-tagged MiD49 (n=6). **(B)** *In vitro* GST pull-down assay. Bacterially expressed PI31-GST and GST control immobilized on a GST column and incubated with MiD49/51 rabbit reticulocyte lysates (top). Coomassie staining of GST-tagged proteins (bottom) (n=3). **(C)** Co-immunoprecipitation of MiD49/51 and Fbxo7 from HEK293T cells transfected with MiD49/51-Myc and FLAG-Fbxo7 constructs. Fbxo7 immunoprecipitated from lysates using anti-FLAG beads (n=2). **(D)** *In vitro* MiD49 ubiquitination assay. MiD49 IVT rabbit reticulocyte lysates incubated with purified SCF Fbxo7 ligase ((WT or ΔF-box) and ubiquitin mix (E1, E2, ubiquitin and ATP) (n=3) **(E)** *In vivo* MiD49 ubiquitination assay. HEK293T cells were co-transfected with ubiquitin-HA, Flag-Fbxo7 (WT or ΔF-box) and MiD49-Myc and MiD49-Myc was immunoprecipitated from lysates. Proteins were resolved via SDS-PAGE and probed with the antibodies indicated (n=3). **(F)** Co-immunoprecipitation of MiD49 and Fbxo7 from HEK293T cells overexpressing MiD49-Myc with WT or pathological mutant variants of Fbxo7 (T22M, L250P, R378G, R498X) (n=3). **(G)** *In vivo* ubiquitination assay. MiD49-Myc was overexpressed in HEK293T cells with FLAG-Fbxo7 (WT, ΔF-box or L250P) and immunoprecipitated from lysates with anti-Myc beads (n=5). **(H)** Immunoblots of endogenous MiD49 and MiD51 protein levels in control and L250P patient fibroblast lysates. Protein levels quantified relative to tubulin loading control (n=3). **(I)** Immunoblots of endogenous MiD49 and MiD51 protein levels in HEK293T cells expressing control and Fbxo7 shRNA. MiD49 and MiD51 protein levels quantified relative to GAPDH and tubulin respectively (n=3). **(J)** HEK293T cells stably expressing control and Fbxo7 shRNA were treated with 250μM H_2_O_2_ for up to 6 hours. Cell lysates were resolved by SDS-PAGE and immunoblotted for MiD49 (n=4). *Error bars = SEM *p<0.05, **p<0.01, ***p<0.001*

### SCF^Fbxo7^ ubiquitinates MiD49

The *in vivo* and *in vitro* interaction data suggest PI31 might recruit MiD49 to the SCF^Fbxo7^ ligase. To test whether Fbxo7 interacted with MiD49 and MiD51, HEK293T cells were co-transfected with FLAG-tagged Fbxo7 and Myc-tagged MiD49/51. Both proteins were detected in anti-FLAG immunoprecipitates of Fbxo7 (Fig. 2C, S1B). Of note, a higher molecular weight smear, indicative of poly-ubiquitination was detected in immunoblots for MiD49-Myc (Fig. 2C), but not MiD51-Myc (Fig. S1B), suggesting MiD49, but not MiD51, is a substrate of SCF^Fbxo7^ ligase. We mapped the Fbxo7-MiD49 interaction using Fbxo7 truncations of functional domains within Fbxo7 in co-IP assays (Fig. S1C). The smallest construct tested, containing only the FP dimerization and F-box domains (aa169-398), interacted with both MiD proteins (Fig. S1D, S1E). As Fbxo7 lacking its F-box domain interacted with both MiD proteins, these data suggest the FP dimerization domain of Fbxo7 mediates interactions with MiD49 and MiD51.

We directly tested the ability of SCF^Fbxo7^ ligase to ubiquitinate MiD49 and MiD51 using *in vitro* ubiquitination assays. Increased higher molecular weight smearing was detected for MiD49 upon addition of full-length Fbox7 but not the ligase-dead ΔF-box (Fig. 2D). No ubiquitination was detected for MiD51 (Fig. S1F). We also conducted *in vivo* ubiquitination assays where HEK293T cells with reduced Fbxo7 levels were co-transfected with Myc-tagged MiD49 or MiD51, FLAG-tagged Fbxo7 constructs, and HA-tagged ubiquitin. Myc-tagged MiD49 or MiD51 was isolated from cells via anti-Myc immunoprecipitation and probed by immunoblotting. MiD49, but not MiD51, showed increased ubiquitination in the presence of the WT Fbxo7 but not the ligase dead ΔF-box allele (Fig. 2E, S1G). Thus, despite both structurally similar MiD proteins interacting with Fbxo7, only MiD49 is ubiquitinated by SCF^Fbxo7^ ligase.

### The Fbxo7-L250P mutation disrupts the ubiquitination of MiD49

We tested whether the Fbxo7-L250P mutation, which removes PI31 interaction, affected Fbxo7 interaction with the MiD proteins. HEK293T cells with constitutive knockdown of Fbxo7 were co-transfected with MiD49 or MiD51 and pathological Fbxo7 mutations (T22M, L250P, R378G and R498X). None of the Fbxo7 mutations affected the interaction with either of the MiD proteins (Fig. 2F, S1H). However, a marked decrease in higher molecular weight species was observed when MiD49 was co-transfected with Fbxo7-L250P compared to WT (Fig. 2G). *In vivo* ubiquitination assays confirmed the reduced ability of SCF^Fbxo7-L250P^ to ubiquitinate MiD49 in cells (Fig. 2G). We demonstrated SCF^Fbxo7-L250P^ is a functional ligase, by verifying its ability to ubiquitinate a known SCF^Fbxo7^ substrate, GSK3β (Fig. S2A); however, its ability to ubiquitinate anther substrate Tomm20 was impaired, both *in vitro* and *in vivo* (Fig. S2B, S2C). These data suggest PI31 is not required for the Fbxo7-MiD49 interaction but is required for SCF^Fbxo7^ ubiquitination of MiD49. Thus, the SCF^Fbxo7-L250P^ ligase has an altered substrate range compared to WT.

### Fbxo7 stabilises MiD49/51 protein levels

We next tested the effect of the L250P mutation on MiD proteins in the patient fibroblasts. Surprisingly, we observed 60% and 65% decreases in MiD49 and MiD51 levels, respectively (Fig. 2H), indicating Fbxo7 and/or PI31 stabilises MiD protein levels. We infer this stabilising effect of Fbxo7 on the MiD proteins is not dependent on its ubiquitination since SCF^Fbxo7-L250P^ does not ubiquitinate MiD49, and MiD51 is not an SCF^Fbxo7^ substrate.

To test whether knock-down of Fbxo7 levels phenocopies the effect of the L250P point mutation, we immunoblotted lysates from HEK293T cells with constitutively expression of a shRNA targeting Fbxo7. We found an approximately 60% decrease in endogenous MiD49 protein levels, indicating Fbxo7 stabilises MiD49 levels (Fig. 2I). A similar effect of reduced Fbxo7 expression was seen on MiD51 levels which were 55% lower. A stabilising effect of Fbxo7 was also seen in cycloheximide chase experiments on cells treated with hydrogen peroxide to induce ROS. Where control cells maintain high levels of Mid49 over 6 hours, cells lacking Fbxo7 show markedly reduced MiD49 levels (Fig. 2J). These data indicate that Mid49/51 levels are stabilized by Fbxo7, under basal and stress conditions.

### Mitochondrial content and function are reduced in L250P fibroblasts

Our data suggest the Fbxo7 L250P mutation might affect the mitochondrial network because of decreased MiD49 and MiD51 protein levels. To test this, patient and control fibroblasts were stained with MitoTracker Green FM, which stains total mitochondrial mass, and mean fluorescence was analysed by flow cytometry. Surprisingly, quantification of MitoTracker Green staining showed an approximately 40% decrease in total mitochondrial content in L250P fibroblasts compared to control cells (Fig. 3A). To confirm this decrease, mitochondrial mass was also measured by qPCR for mtDNA levels. The average copy numbers of three mitochondrial genes (COX I, CYTB and ATP6) were measured and normalised to two regions of genomic DNA (ATP5B and CHR10). Consistent with measurements of mitochondrial staining by FACS analysis, there was an approximately 50-60% decrease in the levels of mtDNA in L250P fibroblasts compared to control cells, indicating patient fibroblasts have a reduced mitochondrial mass (Fig. 3B).

**Figure 3:**
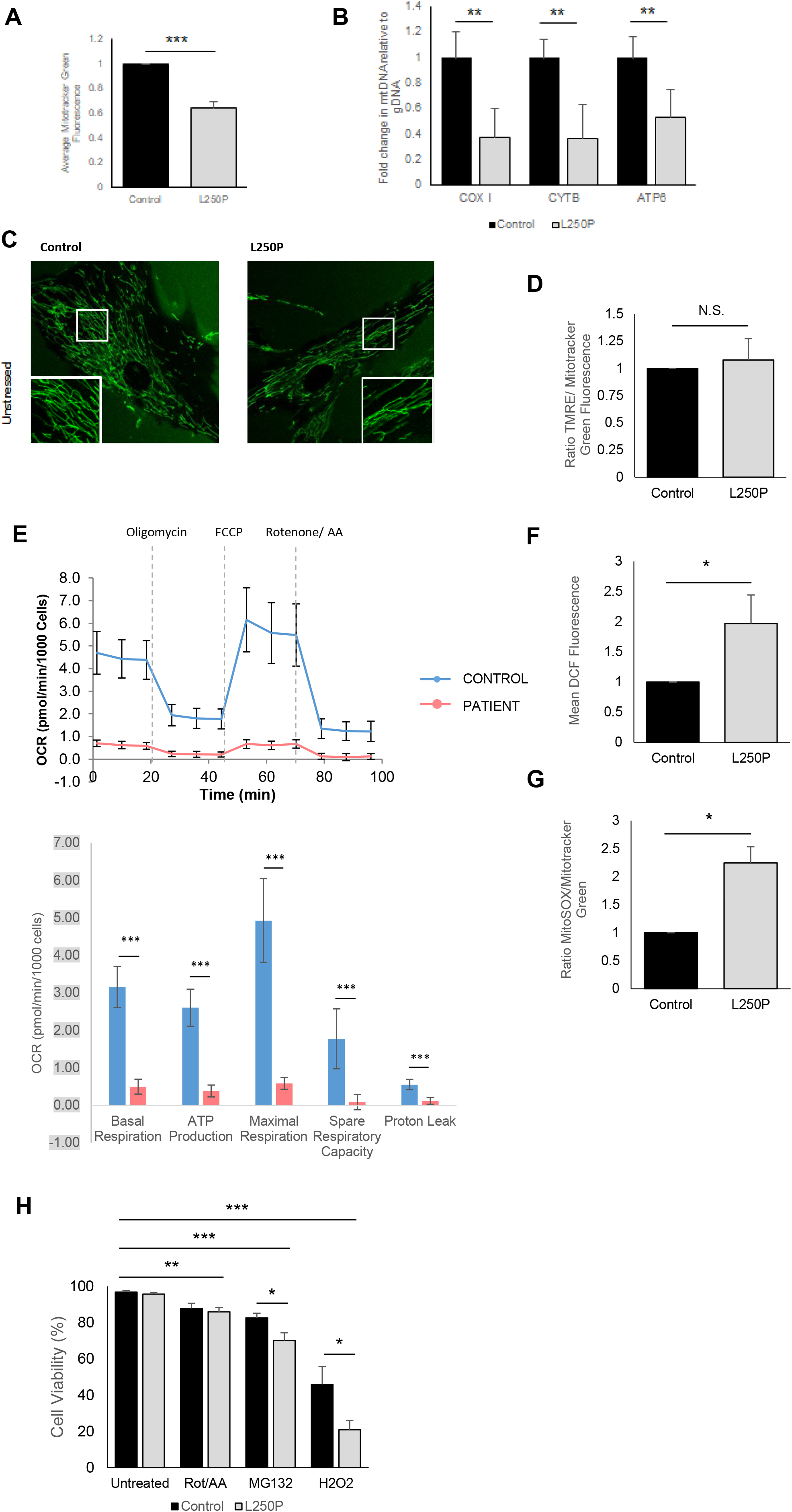
Mitochondrial content and function are reduced in L250P fibroblasts. **(A)** Control and L250P fibroblasts stained with MitoTracker Green and average mitochondrial content was analysed by flow cytometry (n=10). **(B)** Genomic DNA was isolated from control and patient fibroblasts and mtDNA levels were quantified via qPCR. Three mtDNA loci were analysed and normalised to two regions of genomic DNA (n=5). **(C)** Representative images of control and L250P fibroblasts stained with MitoTracker Green and imaged via confocal microscopy. Cells imaged under basal conditions or following treatment with 2-deoxyglucose (n=3). **(D)** Patient and control fibroblasts were incubated with 25nM TMRE and 100nM MitoTracker Green. Mitochondrial membrane potential (as indicated by TMRE fluorescence) was measured via flow cytometry and normalised to mitochondrial content (MitoTracker Green fluorescence) (n=5). **(E)** Representative Seahorse profile and quantification of patient and control fibroblasts analysed by Agilent Seahorse Mito Stress Test (n=3). **(F)** Levels of ROS in L250P and control fibroblasts was measured via flow cytometry. Total cellular ROS was assayed by treating cells with H_2_DCFDA and measuring the resulting DCF fluorescence (n=3). **(G)** Mitochondria specific ROS was assayed by staining cells with MitoSOX. Resulting fluorescent signal was normalised to mitochondrial content (MitoTracker Green) (n=3). **(H)** Control and patient fibroblasts were treated with a panel of cellular stressors and stained with propidium iodide. Cell viability was measured via flow cytometry (n=3). *Error bars = SEM *p<0.05, **p<0.01, ***p<0.001*

We next visualised the mitochondrial network by staining fibroblasts with MitoTracker Green FM and performing confocal microscopy (Fig. 3C). Images were processed using the ImageJ PlugIn MINA, and in agreement with our findings on mass, the overall mitochondrial footprint was significantly lower in the Fbxo7-L250P fibroblasts compared to controls (Fig. S2D). However, no other parameters, including the mean branch length, network size, or the number of individual mitochondria, were significantly altered (Fig. S2D), suggesting that the L250P mutation does not significantly affect mitochondrial network structure/dynamics.

To test if there were any changes in membrane polarisation between the patient and control fibroblasts, we stained cells with TMRE, a fluorescent dye which is only able to enter mitochondria and fluoresce when there is an intact membrane potential. When normalised to overall mitochondrial mass, we did not detect any differences in membrane potential (Fig. 3D)

Given the significant decrease in mitochondrial mass, we evaluated oxidative phosphorylation using a Seahorse XF Cell Mito Stress Test to measure cellular oxygen consumption. Basal respiration, maximal respiration, ATP production and respiratory spare capacity were all found to be significantly lower in the Fbxo7-L250P fibroblasts (Fig. 3E), indicating that the patient cell line harbours a severe respiratory defect.

### Fbxo7-L250P patient fibroblasts show increased levels of ROS

Deficiencies in Fbxo7 have previously been shown to be associated with increases in the level of cytosolic ROS (23, 43, 44). To investigate if patient fibroblasts also had changes in ROS levels, we treated cells with the ROS indicator H_2_DCFDA. Non-fluorescent H_2_DCFDA is converted to fluorescent DCF following oxidation by ROS; thus, increased fluorescence is indicative of higher levels of oxidative stress. Consistent with previous findings on Fbxo7 reducing ROS levels, patient fibroblasts had two-fold higher levels of cellular ROS compared to control fibroblasts (Fig. 3F).

To assay for mitochondrial ROS, cells were incubated with MitoSOX, a derivative of dihydroethidium, which accumulates in mitochondria, and upon oxidation by superoxide, produces a fluorescent signal. No significant differences in total mitochondrial ROS levels were detected, suggesting mitochondrial ROS is not likely to account for the two-fold increase in cellular ROS detected with H_2_DCFDA (Fig. 3F). However, when normalised to mitochondrial mass, the MitoSOX signal in L250P fibroblasts was approximately double that of control cells (Fig. 3G). These data indicate that patient fibroblasts have elevated levels of mitochondrial and cellular ROS, although, we did not detect any differences in cell viability as a result of this increase in ROS between the control and patient fibroblasts under basal conditions. However, the patient fibroblasts were found to be less viable following treatment with cellular stressors including hydrogen peroxide and the proteasome inhibitor MG132, but not to rotenone and antimycin A (Fig. 3H). Consistent with the reduced proteasome activity and higher intracellular ROS in the patient cells, these data suggest that the patient cells are more sensitive to proteasome inhibition and increased exposure to hydrogen peroxide than control cells, but both cell lines respond equivalently to mitochondrial inhibition.

## Discussion

In this study, we report on a patient who was diagnosed at 5 months of age with clinical features suggestive of a complicated pyramidal syndrome. Whole exome sequencing revealed a new homozygous point mutation, c.749T>C in *FBXO7* (p.Leu250Pro, NM_012179.4), which has not been reported previously. Bi-allelic loss of function mutations in *FBXO7* are associated with an early onset parkinsonian-pyramidal syndrome termed PARK15 (OMIM# 260300) typically presenting during the juvenile period (4, 6). Our patient presented an unusual and to our knowledge, the most severe phenotype with infantile onset and significant neurological impairment with minimal L-dopa responsiveness and cognitive impairments. Degenerating axons in the optic nerves have been described in conditional mouse lacking Fbxo7 specifically in the myelinating cells of the central and peripheral nervous system (44), highlighting an important role for Fbxo7 in maintaining the optic nerve fibres. At the age of 2.5 years, an evolving optic atrophy has also been detected in this patient, suggesting that this new point mutation could have an effect on Fbxo7 functions in the optic nerve axons. Moreover, in other mouse models lacking Fbxo7 specifically in the dopaminergic neurons, decreased numbers of fibres innervating the striatum are observed (28, 34), suggesting that Fbxo7 could have a more global role in axonal growth in different parts of the nervous system, and this would explain the abnormalities on the DAT SPECT scan.

The L250P mutation lies within the FP domain of Fbxo7, and it was previously reported that mutation of V253E prevents Fbxo7 interaction with PI31 (26). Consistent with these previous studies on the mode of interaction of the FP domains of Fbxo7 and PI31, the L250P mutation specifically abolishes their hetero-dimerization. In characterising the fibroblasts from this patient, we have shown the levels and stability of Fbxo7-L250P and PI31 are decreased, highlighting the importance of the β-interface for the interaction between the two proteins. Of note, the L250P mutation did not prevent homo-dimerization, suggesting other regions of Fbxo7 mediate this and that homo-dimerization is not sufficient to stabilize Fbxo7-L250P protein levels.

Multiple studies have reported Fbxo7 and PI31 interact with, and affect, the proteasome, ranging from assembly and activity to trafficking (27, 34, 35). Our studies in patient fibroblasts adds to this picture showing reduced Fbxo7 and PI31 protein levels and the loss of their heterodimerization correlated with lower levels of specific proteasome subunits, including an immunoproteasome subunit, and significantly reduced proteasome activity. These observations underscore the multiple ways in which Fbxo7 and PI31 can affect the proteasome.

Given that PI31 is usually thought of as a proteasome regulator, a surprising finding from our studies is that PI31 affects the substrate range of the SCF^Fbxo7^ E3 ligase. Since we showed mutant Fbxo7-L250P could bind to Skp1, we reasoned it could be incorporated into an active E3 ubiquitin ligase. We discovered the MiD proteins as direct interactors of PI31, via its FP domain, and showed MiD49 is a PI31-dependent substrate of SCF^Fbxo7^ ligase. In principle, other interacting proteins of PI31 could similarly be recruited for ubiquitination. PI31 appears to modulate the substrate range as evidenced by SCF^Fbxo7-L250P^ ligase ubiquitinating known Fbxo7 substrates like GSK3β, but not others, like Tomm20 and MiD49. Interestingly, MiD49 still interacted with Fbxo7-L250P but was not ubiquitinated, suggesting PI31 binding promotes a conformational change that permits MiD49 ubiquitination. These data argue PI31 acts as a co-factor for SCF^Fbxo7^ facilitating ubiquitination of a subset of its substrates, and the L250P mutation is a change-of-function mutation which alters the Fbxo7 substrate repertoire.

We found the levels of MiD49 and MiD51 protein were significantly reduced in the patient fibroblasts, suggesting Fbxo7 and/or PI31 stabilise their levels. MiD49 and MiD51 are adaptor proteins which recruit Drp1 to catalyse the fission of organelles, including mitochondria, ER, and peroxisomes. Although some studies report loss of MiD49 and MiD51 levels can cause an elongated, hyper-fused mitochondrial network (43, 45, 46), this was not the case in the patient fibroblasts. Since the MiD proteins function in a redundant fashion with other adaptors Fis1 and Mff, it is possible that these proteins compensated for the decreased expression of the MiD proteins in these cells. The presence of multiple Drp1 adaptors suggests cell type-specific or contextual roles for these proteins (42, 47, 48). More recent studies support this idea, e.g., MiD49 has been shown to be involved in localised mitochondrial fission caused by damage to the plasma membrane, while MiD51 is important for coordinating the Bcl2 family of proteins on mitochondria in response to apoptotic stimuli (49, 50). Thus, the phenotypes of MiD49 and MiD51 loss may be responsive to the physiological state of the cell. Despite the lack of an effect on the mitochondrial network, the L250P fibroblasts showed a significant reduction in mitochondrial mass and in oxidative phosphorylation, and higher levels of cytosolic and mitochondrial ROS. Impaired mitochondrial respiration and higher levels of cytosolic ROS have been reported in another patient fibroblast line bearing homozygous R378G mutations in Fbxo7, although it is not known whether there are effects on the proteasome in these cells (23). The involvement of Fbxo7 in regulating proteasomes and mitochondria involves two of the commonly dysregulated pathways associated with neurological diseases. Our data implicate PI31 as a coregulating factor with Fbxo7 that impacts on both of these pathways.

## Materials & Methods

### Ethical approval

Informed consent for participation in the study was obtained from the parents of the patient. The study was approved by the Emek Medical Center ethics committee (study no. EMC-0067-09).

### Whole exome sequencing

Exonic sequences were enriched with the SureSelect Human All Exon 50□Mb V5 Kit (Agilent Technologies, Santa Clara, CA, USA). Sequences were generated on a HiSeq2500 (Illumina, San Diego, CA, USA) as 125-bp paired-end runs. Read alignment and variant calling, were performed with DNAnexus (Palo Alto, California, USA) using default parameters with the human genome assembly hg19 (GRCh37) as reference. Exome analysis of the proband yielded 55 million reads, with a mean coverage of 97X. Following alignment to the reference genome [hg19] and variant calling, variants were filtered out if the total read depth was less than 8X and if they were off-target (>8 bp from splice junction), synonymous, or had minor allele frequency (MAF)□>□0.005 in the gnomAD database. Homozygous variants of interest that survived filtering are provided in Table S1.

### Cell lines

HEK923T and HFF-1 control fibroblasts (Human Foreskin Fibroblasts) were purchased from ATCC. Cells were maintained in DMEM supplemented with 10% heat-inactivated foetal bovine serum (Gibco), 100 U/mL penicillin and streptomycin (Gibco) at 37°C in a humidified 5% CO_2_ atmosphere. Cell lines stably expressing shRNA to human *FBXO7* were generated as described previously (20) and selected with 2 μg/mL puromycin (Sigma-Aldrich).

### Antibodies

Antibodies against the following proteins were used for immunoblotting: Fbxo7 (generated in (21)), Fbxo7 (Aviva ARP43128_P050), Skp1 (BD Biosciences 610530), Myc-tag (CST 2272), FLAG-tag (Sigma F3165), Tubulin (Sigma T6557), GAPDH (Sigma G9545), PI31 (Enzo PW9710-0025), HA-tag (Cell Signalling 3724S), LMP7 (Abcam 3329), PSMA7 (Genetex GTX101745), PSMC4 (Santa Cruz sc-166115), PSMB1 (Biorbyt orb39549), MiD51 (Santa Cruz, sc-514135), MiD49 (ab101350). Signal detection was enhanced by chemiluminescence (ECL) (GE Healthcare) or SuperSignal^™^ West Pico PLUS Chemiluminescent Substrate (Thermo Scientific).

### DNA constructs

pcDNA3 vectors expressing full length, truncated or ΔF-box Fbxo7 with an N-terminal Flag tag or C-terminal HA tag have been previously described (21, 24). Flag-Fbxo7 L250P construct in pcDNA3 has been engineered by site-directed mutagenesis and cloned into Flag-pcDNA3 vector. Fbxo7 L250P-HA construct was generated by subcloning Fbxo7-L250P from the Flag-pcDNA3 vector into pcDNA3-HA vector.

### Cell transfections

HEK293T cells were split 24 hours prior transfection when cells were at 50 to 80% confluency. For each 10cm dish, 2μg to 6μg of plasmid DNA was diluted in 300ul of Optimem media (Invitrogen) and mixed with the transfection reagent (Fugene or PEI) at a ratio of 3μg/μl and incubated at RT for 15 minutes before adding to cells. Cells were transfected for 36-48 hours prior to harvesting.

### Immunoprecipitation

For immunoprecipitation, cells were lysed in RIPA Lysis Buffer (50 mM Tris-HCl (pH 7.6), 150mM NaCl, 1% NP-40, 0.1% SDS, 0.5% sodium deoxycholate) or 0.1% NP-40 lysis buffer with a protease inhibitor cocktail (Sigma-Aldrich) and other inhibitors (1 mM PMSF, 10 mM NaF, 1 mM Na_3_VO_4_).

Fbxo7 was immunoprecipitated from cell lysates using 2μl of anti-Fbxo7 antibody (Aviva), or isotype matched control, for 1 hour at 4°C with rotation, then 20 μL Protein A/G PLUS-Agarose (Santa Cruz) was added and samples were incubated for a further 3 hours. Beads were washed 4 times in lysis buffer and resuspended in 2X Laemmli loading buffer.

For co-immunoprecipitation assays, cells were lysed as above. An anti-FLAG^®^ M2 Affinity Gel (Sigma-Aldrich) was washed twice with lysis buffer then added to lysates and incubated for 4 hours at 4°C with rotation. Beads were washed 4 times in lysis buffer and beads were resuspended in 30μl of 2X Laemmli loading buffer.

### Purification of SCF^Fbxo7^ complexes and substrates

HEK293T cells were transfected with Skp1, Cullin1 and Myc-Rbx1, alongside FLAG-Fbxo7 constructs. After 48 hours, cells were resuspended in KCl lysis buffer (50 mM Tris-HCl pH 7.5, 225 mM KCl, 1% NP-40) with a protease inhibitor cocktail (Sigma-Aldrich) and other inhibitors (1 mM PMSF, 10 mM NaF, 1 mM Na_3_VO_4_). Lysates were incubated with Anti-FLAG^®^ M2 Affinity Gel (Sigma-Aldrich) for 4 hours at 4°C with rotation. Beads were washed 3 times in lysis buffer and twice in elution buffer (10 mM HEPES, 225 mM KCl, 1.5 mM MgCl_2_, 0.1% NP-40), and protein was eluted with 100 μg/mL FLAG peptide (Sigma-Aldrich) in elution buffer for 1 hour at 4°C with rotation. Purified SCF complexes were stored at −20°C in 15% glycerol.

### In vitro *ubiquitination assays*

To purify HA-tagged substrates for *in vitro* ubiquitination, HEK293T cells were transfected with HA-Tomm20 or HA-GSK3β which was immunoprecipitated with anti-HA agarose (Sigma-Aldrich) 48 hours after transfection. Substrate was eluted with 300 μg/mL HA peptide (Sigma-Aldrich) and stored at 20°C in 15% glycerol. Myc-tagged MiD49 and MiD51 were produced via *in vitro* transcription/translation (IVT) (Promega). For *in vitro* ubiquitination assays, a ubiquitin mix was prepared with 100 nM E1 (UBE1, Bio-Techne), 500 nM E2 (UbcH5a, Bio-Techne), 20 μM human recombinant ubiquitin (Santa Cruz) and 2 mM ATP (Bio-Techne) in 1X ubiquitin conjugation reaction buffer (Bio-Techne). This was incubated for 5 min at RT then added to 100 nM SCF and 1 μL substrate and incubated for 1 hour at 30°C. The entire 10 μL reaction was mixed with an equal volume of 2X Laemmli loading buffer, resolved by SDS-PAGE and analysed by immunoblotting.

### *In vivo* ubiquitination assays

HEK293T cells were transfected with tagged ubiquitin, FLAG-Fbxo7 and substrate constructs. Cells were treated with 25 μM MG132 for 4 hours before being harvested and lysed in RIPA lysis buffer.

For MiD49/51 ubiquitination assays, lysates were incubated with 2 μl anti-Myc-tag antibody for 1 hour at 4°C with rotation. 20 μl Protein A/G PLUS-Agarose (Santa Cruz) was then added to samples and incubated for a further 2 hours. For Tomm20 ubiquitination assays, 20 μl anti-HA agarose (Sigma-Aldrich) was added to lysates for 3 hours at 4°C with rotation. Beads were washed 4 times in lysis buffer and resuspended in 2XLaemmli loading buffer.

### GST protein purification

GST-tagged protein constructs were transformed into FB810 *E. coli*. Bacterial cultures were grown in 2xTY medium containing 100μg/ml ampicillin at 37°C until an optical density of between 0.4-0.6 was reached. Protein expression was induced with 1mM IPTG and cells were incubated at 30°C for 3 hours. Cells were pelleted by centrifugation (3000 x g, 15 mins, 4°C) then resuspended in 4ml TBS buffer with cOmplete, EDTA-free protease inhibitor cocktail (Roche) and lysozyme (Sigma-Aldrich). Cell suspensions were disrupted by sonication (3x 20 seconds) on a SoniPrep 150. Triton X-100 in TBS was added to a final concentration of 1% and samples were centrifuged at 18,000 x g, 25 mins, 4°C. Glutathione Sepharose 4B beads (GE Healthcare) were added to supernatants and rotated at 4°C for 2 hours to capture GST-tagged proteins. Beads were washed in 0.5% Triton X-100. Glycerol was added to a final concentration of 15% and stored at −20°C.

### GST Pull-down assay

MiD49 and MiD51 were produced via IVT and diluted in Alternative binding buffer (20mM Hepes-KOH pH7.6, 50mM KCl, 2.5mM MgCl2, 10% glycerol, 0.02% NP-40, 1mM PMSF, 1mM DTT). Diluted lysates were added to equal amounts of GST proteins bound to beads and rotated for 2 hours at 4°C. Beads were washed 5 times in NET-N wash buffer (50mM Tris pH7.5, 1mM EDTA, 1% NP-40, 150mM NaCl) and resuspended in 2xLaemmli loading buffer.

### Proteasome assay

Proteasome-Glo Chymotrypsin-like assay (Promega G8621) was used for proteasome assays. One confluent 10cm dish of cells was washed in PBS before trypsinization. The collected cells were washed twice in culture media to remove trypsin. Cells were lysed in a lysis buffer (10mM Tris-HCl, 10mM NaCl, 2mM EDTA, 0.5% Triton). 50μg of protein lysates in 50μl total volume was used to mix with 50μl assay reagent. Control samples were treated with the proteasome inhibitor MG132. Samples were loaded in triplicates into a white-walled multi-well plate. After 30 minutes of incubation, plates were read in a luminometer.

### Cycloheximide (CHX) chase assay

Cells were seeded equally onto a 6-well plate. At around 80% confluence, the first well was harvested, and the rest of the cells were treated with 100ug/mL of CHX. Each subsequent well was harvested at timepoints of 2, 4-, 6-, 8- and 12-hours post-treatment, and frozen on dry ice. Cells were stored at −80°C until time for lysis in 50μl RIPA lysis buffer. Samples were resolved by SDS-PAGE and immunoblotted.

### mtDNA quantification

Genomic DNA was harvested from cells (Wizard Genomic DNA Isolation Kit). qPCR reactions were performed in triplicate using SYBR^®^ Green JumpStart^™^ Taq ReadyMix^™^ (Sigma). The copy number of three mitochondrial genes COX1 (Cytochrome C oxidase 1), CYTB (Cytochrome B) and ATP6 (ATP synthase 6) was normalized against the copy number of the nuclear gene ATP5B (ATP synthase subunit beta) and a nuclear gene desert region of chromosome 10 to calculate the relative mtDNA copy number (oligonucleotide sequences from (51)).

### Seahorse Mito stress test

The Seahorse Cell Mito Stress Test was performed according to the manufacturer’s instructions. In brief, the day prior to the assay, cells were harvested and plated into a Seahorse XF96 Cell Culture Microplate (Agilent) at 1000 cells/well and incubated at 37°C overnight. On the day of the assay, standard cell culture media was replaced with Seahorse XF DMEM medium (Agilent), supplemented with 2 mM glutamine (Thermo Scientific), 10 mM glucose (Thermo Scientific) and 1 mM pyruvate (Thermo Scientific), at pH 7.4 and cells were incubated at 37°C for 1 hour in a non-CO2 incubator. Mitochondrial oxygen consumption rate was analysed using the Seahorse XF Cell Mito Stress Test Kit (Agilent) with 1μM oligomycin, 1μM FCCP and 1μM rotenone/antimycin a. The assay was run at 37°C on a Seahorse XF96 analyser (Agilent). OCR measurements were normalised to cell number which was quantified using the CyQUANT^™^ Cell Proliferation Assay Kit (ThermoFisher Scientific). Data was analysed using Wave software (Agilent).

### DCFDA staining for total cellular ROS

Equal numbers of cells were harvested, washed and resuspended in OptiMEM and incubated with 2.5μM H_2_DCFDA for 30 minutes at 37°C. After 30 minutes, cells were washed and resuspended in PBS and propidium iodide was added to a final concentration of 0.4μg/ml. Mean DCF fluorescence in live cells was measured via flow cytometry.

### MitoSOX assay

Equal numbers of cells were harvested, washed, and resuspended in HBSS and incubated with either 100nM MitoTracker Green FM (ThermoFisher Scientific) for 30 minutes or 2.5μM MitoSOX (ThermoFisher Scientific) for 15 minutes at 37°C. Cells were washed and resuspended in PBS and mean fluorescence was measured via flow cytometry. MitoSOX fluorescence was normalised to mitochondrial content.

### Mitochondrial membrane potential assay

Equal numbers of cells were harvested and incubated with 100nM MitoTracker Green FM in DMEM for 30 minutes at 37°C. Cells were washed with PBS and incubated with 25nM TMRE (Abcam) in DMEM for 15 minutes at 37°C. Mean fluorescence was measured via flow cytometry. TMRE fluorescence was normalised to mitochondrial content.

### Cell viability assay

Control and patient fibroblasts were incubated with a range of cellular stressors for 3 hours (1mM hydrogen peroxide) or 24 hours (1μM rotenone/antimycin a, 10 μM MG132). Cells were harvested and washed in PBS. Cells were resuspended in PBS and propidium iodide (PI) was added at 0.4μg/ml. The percentage of PI positive cells was analysed via flow cytometry.

### Mitochondrial Network Analysis

Cells were plated on 35mm glass bottom dishes (Matek) and stained with MitoTracker Green FM for 30 minutes at 37°C. Cells were washed with PBS and fresh media was added. Cells were imaged using an LSM700 confocal microscope (63x magnification). Parameters of mitochondrial network morphology were quantified using the Image J PlugIn MiNA (Mitochondrial Network Analysis).

### Quantification and statistical analysis

Immunoblot analyses were quantified using ImageJ processing software. Data are presented as mean ± SEM. Statistical differences were calculated using Student’s two-tailed t tests with a significant cutoff of p < 0.05.

## Acknowledgements

We thank members of the Laman Lab for helpful discussions. This work is supported by funding from the BBSRC DTP to LS and HL and a Parkinson’s UK grant G-1701 to RAB, MZ and HL. This research is also supported by the NIHR Cambridge Biomedical Research Centre (BRC-1215-20014, including the Cell Phenotyping Hub. The views expressed are those of the author(s) and not necessarily those of the NIHR or the Department of Health and Social Care. This research was funded in whole, or in part, by the Wellcome Trust 203151/Z/16/Z. NQM is supported by the Francis Crick Institute, which receives its core funding (FC001115) from Cancer Research UK, the UK Medical Research Council and the Wellcome Trust. For the purpose of Open Access, the author has applied a CC BY public copyright licence to any Author Accepted Manuscript version arising from this submission.

## Author Contributions

All authors have read and approved the manuscript. SA and LS carried out experiments, analysed data and wrote the manuscript. NQM did structural predictions. VC, OE and RS identified the L250P mutation, conceived of the study, provided patient fibroblasts and patient care. RAB and MZ secured funding and analysed data. HL conceived of the study, analysed data, and wrote the manuscript.

## The Paper Explained

### Problem

The discovery of monogenetic mutations associated with early onset PD allows researchers to focus in on specific pathways to understand the causes of genetic and sporadic cases of PD. It is apparent that several of the genetic forms of PD damage multiple downstream physiological processes, including mitochondria, protein homeostasis and autophagy, stress responses, calcium signalling, metabolism, and intracellular trafficking. This makes it difficult to understand how dysfunction in any particular process affects others and how they link together to cause disease.

### Results

We identified a novel missense mutation in the dimerising domain of the ubiquitin ligase, Fbxo7, which prevents is interaction with a protein, PI31. PI31 is a regulator of the proteasome, and we find patient cells have reduced proteasome activity. Unexpectedly, we find this missense mutation also changes the ability of this ubiquitin ligase to modify a class of proteins involved in controlling the mitochondrial network. The patient cells also have reduced levels of mitochondria and are more vulnerable to cellular stresses. Thus, we show Fbxo7 and PI31 reciprocally regulate each other’s functions.

### Impact

By understanding how downstream physiological processes are co-ordinately affected by pathological mutations occurring in genetic cases of parkinsonism, we can decide which are the most proximal and actionable pathways for designing therapeutics.

---

**Supplementary Figure 1:**
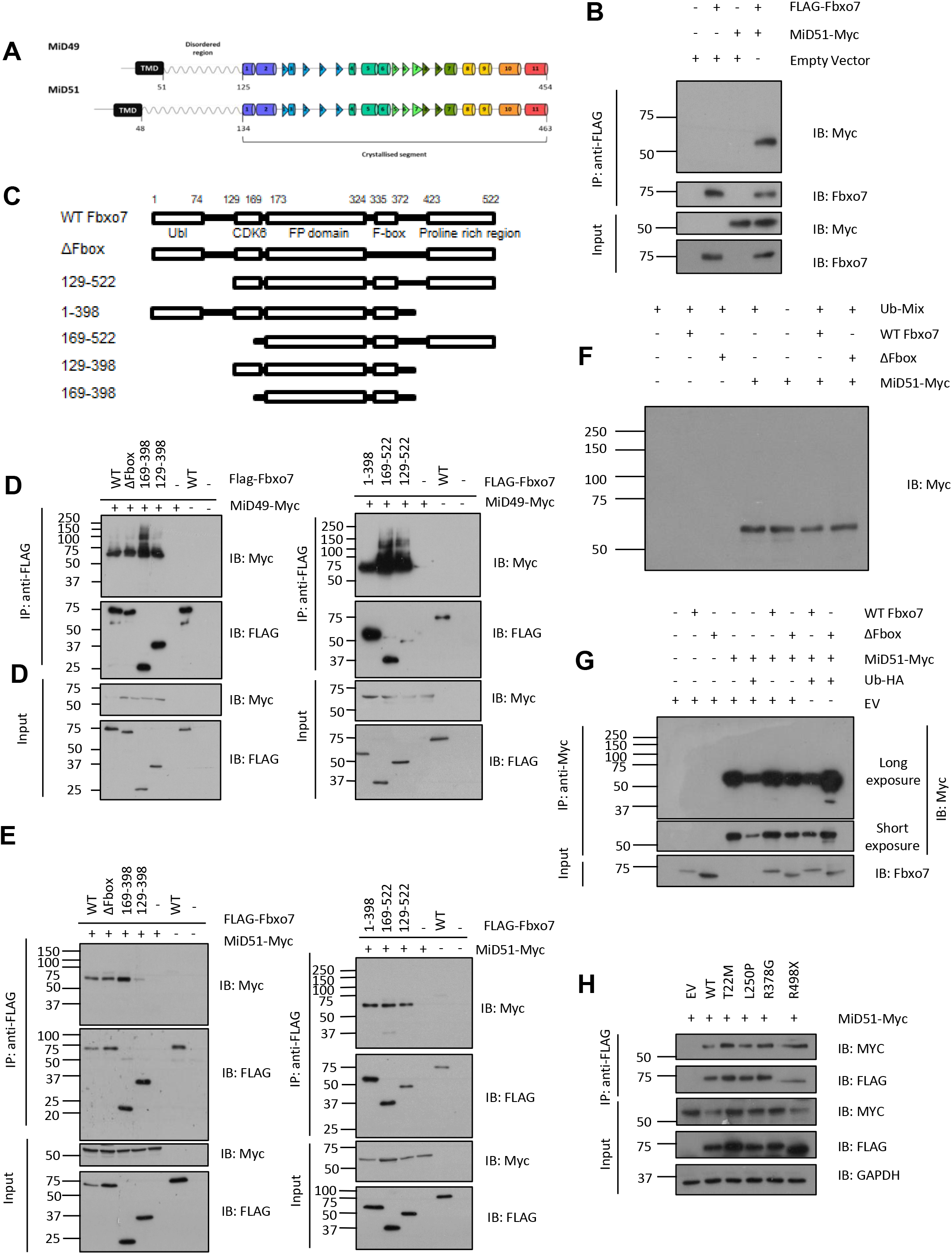
**(A)** Schematic of MiD49 and MiD51 protein structure (wavy line indicates disordered region, cylinders represent α-helices, triangles represent β-strands). (Adapted from (52)). **(B)** Co-immunoprecipitation of MiD51 and Fbxo7 from HEK293T cells transfected with MiD51-Myc and FLAG-Fbxo7. Fbxo7 immunoprecipitated from lysates using anti-FLAG beads (n=2). **(C)** Graphical representation of Fbxo7 N- and C-terminal deletion constructs used for interaction mapping. N-terminal FLAG-tags not shown. **(D, E)** Fbxo7-MiD protein interaction mapping. HEK293T cells co-transfected with MiD49/51 and various Fbxo7 constructs lacking N- and/or C-terminal domains (illustrated in (**C**)). Fbxo7 was immunoprecipitated from lysates with anti-FLAG beads. Proteins were resolved by SDS-PAGE and immunoblots were probed with the antibodies indicated (n=2). **(F)** MiD51 *In vitro* ubiquitination assay. MiD51 IVT rabbit reticulocyte lysates were incubated with purified SCF^Fbxo7^ ligase (WT or ΔF-box) and ubiquitin mix (E1, E2, ubiquitin and ATP) (n=3). **(G)** *In vivo* MiD51 ubiquitination assay. MiD51-Myc immunoprecipitated from HEK293T cells cotransfected with ubiquitin-HA and FLAG-Fbxo7 (WT or ΔF-box) and MiD51-Myc. Proteins were resolved via SDS-PAGE and membranes probed with the antibodies indicated (n=3). **(H)** Co-immunoprecipitation of MiD51 with Fbxo7 from HEK293T cell lysates overexpressing WT Fbxo7 and pathological mutants (T22M, L250P, R378G, R498X) (n=1).

**Supplementary Figure 2:**
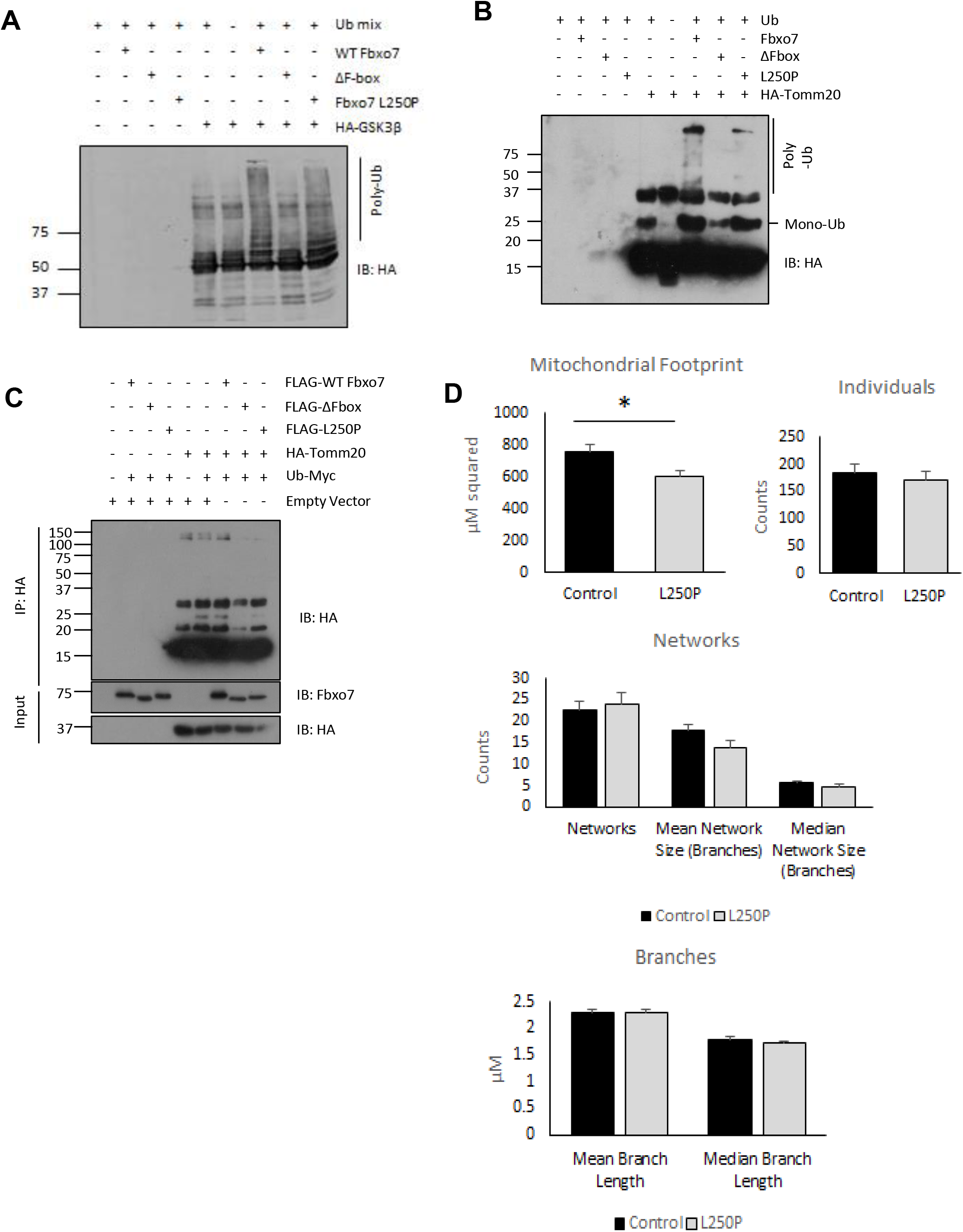
**(A, B)** GSK3β and Tomm20 *In vitro* ubiquitination assay. Purified Fbxo7 (WT, ΔF-box or L250P) SCF complexes were incubated with purified HA-tagged GSK3β (n=2) or Tomm20 (n=2). Proteins were resolved via SDS-PAGE and immunoblotted with the antibodies indicated. **(C)** *In vivo* Tomm20 ubiquitination assay. HA-tagged Tomm20 was overexpressed in HEK293T cells FLAG-Fbxo7 (WT, ΔF-box or L250P). Tomm20 was immunoprecipitated from lysates with anti-HA beads and proteins were resolved via SDS-PAGE gel (n=3) **(D)** Control and L250P fibroblasts stained were MitoTracker Green and imaged via confocal microscopy. Parameters of mitochondrial network morphology were quantified using the MINA Image J PlugIn. Approximately 20-30 images were analysed per repeat for each cell line (n=3). *Error bars = SEM *p<0.05, **p<0.01, ***p<0.001*

